# Implementing high-throughput insect barcoding in microbiome studies: impact of non-destructive DNA extraction on microbiome reconstruction

**DOI:** 10.1101/2024.04.30.591865

**Authors:** Veronika Andriienko, Mateusz Buczek, Rudolf Meier, Amrita Srivathsan, Piotr Łukasik, Michał R. Kolasa

## Abstract

**Background:** Symbiotic relationships with diverse microorganisms are crucial for many aspects of insect biology. However, while our understanding of insect taxonomic diversity and the distribution of insect species in natural communities is limited, we know much less about their microbiota. In the era of rapid biodiversity declines, as researchers increasingly turn towards DNA-based monitoring, developing and broadly implementing approaches for high-throughput and cost-effective characterization of both insect and insect-associated microbial diversity is essential. We need to verify whether approaches such as high-throughput barcoding, a powerful tool for identifying wild insects, would permit subsequent microbiota reconstruction in these specimens.

**Methods:** High-throughput barcoding (“megabarcoding”) methods often rely on non-destructive approaches for obtaining template DNA for PCR amplification by leaching DNA out of insect specimens using alkaline buffers such as HotSHOT. This study investigated the impact of HotSHOT on microbial abundance estimates and the reconstructed bacterial community profiles. We addressed this question by comparing quantitative 16S rRNA amplicon sequencing data for HotSHOT-treated or untreated specimens of 16 insect species representing six orders and selected based on the expectation of limited variation among individuals.

**Results:** We find that in 13 species, the treatment significantly reduced microbial abundance estimates, corresponding to an estimated 15-fold decrease in amplifiable 16S rRNA template on average. On the other hand, HotSHOT pre-treatment had a limited effect on microbial community composition. The reconstructed presence of abundant bacteria with known significant effects was not affected. On the other hand, we observed changes in the presence of low-abundance microbes, those close to the reliable detection threshold. Alpha and beta diversity analyses showed compositional differences in only a few species.

**Conclusion:** Our results indicate that HotSHOT pre-treated specimens remain suitable for microbial community composition reconstruction, even if abundance may be hard to estimate. These results indicate that we can cost-effectively combine barcoding with the study of microbiota across wild insect communities. Thus, the voucher specimens obtained using megabarcoding studies targeted at characterizing insect communities can be used for microbiome characterizations. This can substantially aid in speeding up the accumulation of knowledge on the microbiomes of abundant and hyperdiverse insect species.

## Introduction

Insects have achieved tremendous evolutionary success, reflected in species diversity estimated in millions, functional diversity, and distribution across almost all terrestrial ecosystems, where they fulfill multiple critically important roles (Losey & Vaughan, 2006; Weisser & Siemann, 2008). However, their biodiversity is now in steep decline, with habitat degradation, environmental pollution, and climate change identified as some of the key drivers of biomass and diversity losses estimated at 9% per decade and potentially 40% of all species in the near future (Sánchez-Bayo & Wyckhuys, 2019; Van Klink et al., 2020). Simultaneously, only a fraction, perhaps one-fifth, of insect species are known to science (Stork, 2018).

The dire need to intensify insect biodiversity characterization and monitoring efforts has not been missed by the scientific community, with the rapid development of approaches such as metabarcoding or high-throughput cost-effective individual barcoding (Srivathsan et al., 2024), enabling the characterization of entire insect community samples (Iwaszkiewicz-Eggebrecht et al., 2023). On the other hand, the description of approaches for the study of shifting biotic interactions within the monitored multi-species communities has lagged. Arguably, the most significant of these associations, yet poorly known outside of a limited set of model species, are those with symbiotic microorganisms: bacteria and fungi that inhabit insect bodies and can dramatically affect their ecology and evolution (McFall-Ngai et al., 2013; Łukasik & Kolasa, 2023).

The diversity of insects is reflected in at least a comparable diversity of microbial symbionts. Different functional categories of symbionts have diverse and often pivotal effects on the life history traits of their insect hosts, influencing their biology in various ways (Łukasik & Kolasa, 2023). Particularly well-known are the microbial roles in the biology and evolution of clades that feed on nutrient-limited diets, including sap-feeding hemipteran clade Auchenorrhyncha (Moran, McCutcheon & Nakabachi, 2008) or blood-feeding insects like bedbugs (Husnik, 2018). On the other hand, through their effects on traits such as reproduction and defense against biotic and abiotic factors (Lemoine, Engl & Kaltenpoth, 2020), symbionts can strongly influence the biology of insects on much shorter timescales, providing selective advantages to hosts and thus shaping their population dynamics and ecological interactions (Ferrari & Vavre, 2011; Łukasik & Kolasa, 2023). Some insect groups, including diverse ants (Sanders et al., 2017), have not developed specific symbiotic relationships but may still harbor transient microbes derived from food or other environmental sources, yet their overall abundance may be low (Hammer, Sanders & Fierer, 2019). Moreover, pathogens can dramatically affect individuals and populations, whether they target insects or are vectored by them (Dwyer, Dushoff & Yee, 2004). We know little about the distributions of different functional categories of these microbes within and across multi-species natural communities. At the same time, insect communities themselves remain poorly characterized in various ecosystems. To address the gaps in our understanding of insect diversity, several initiatives have embarked on high-throughput barcoding (“megabarcoding”) (Chua et al., 2023) and metabarcoding in the past decade (Meier et al., 2016; Geiger et al., 2016; Buchner et al., 2024). These are improving our understanding of insect community compositions (Srivathsan et al., 2023), and efforts have been made to significantly reduce costs in order to conduct such surveys at a large scale effectively (Meier et al., 2016; Srivathsan et al., 2021). To go beyond characterizing insect communities to addressing questions about insect microbial diversity, distribution, and roles across large multi-species insect collections, whether representing particular clades or community samples, we need robust, cost-effective workflows that simultaneously provide insect identity information with an unbiased picture of their microbiota.

Most studies on insect microbiomes examine target species of interest but have not looked at them in terms of insect communities (Kolasa et al., 2019; Nakabachi, Inoue & Hirose, 2022). Some authors attempted surveying microbiota using the whole bulk multi-species samples, the same as those used for insect metabarcoding (Gibson et al., 2014). This is, however, limiting given that it dissociates the microbial data from specimen identity. The alternative strategy comprises processing insect specimens individually using megabarcoding (Srivathsan et al., 2023). Recent megabarcoding approaches aim to obtain reference DNA barcodes for insect specimens using non-destructive DNA extraction approaches that give sufficient template DNA for PCR while preserving morphological features and most of the genomic DNA within the specimen (Srivathsan et al., 2024). This is usually achieved by treating the specimen with alkaline buffers to leach the DNA out of the specimen (Fig. 1B), which can then be used as a template for PCR. DNA barcoding then involves PCR amplification and sequencing, and the resulting barcode information can be used to group the physical specimens into mOTUs (molecular Operational Taxonomic Units) for further morphological examinations. These specimens can also be used to obtain associated microbial data and assess them using approaches such as marker gene amplicon or metagenome sequencing. 16S rRNA amplicon sequencing has been a popular technique for characterizing microbial community composition across multiple samples (Langille et al., 2013).

**Figure 1.**
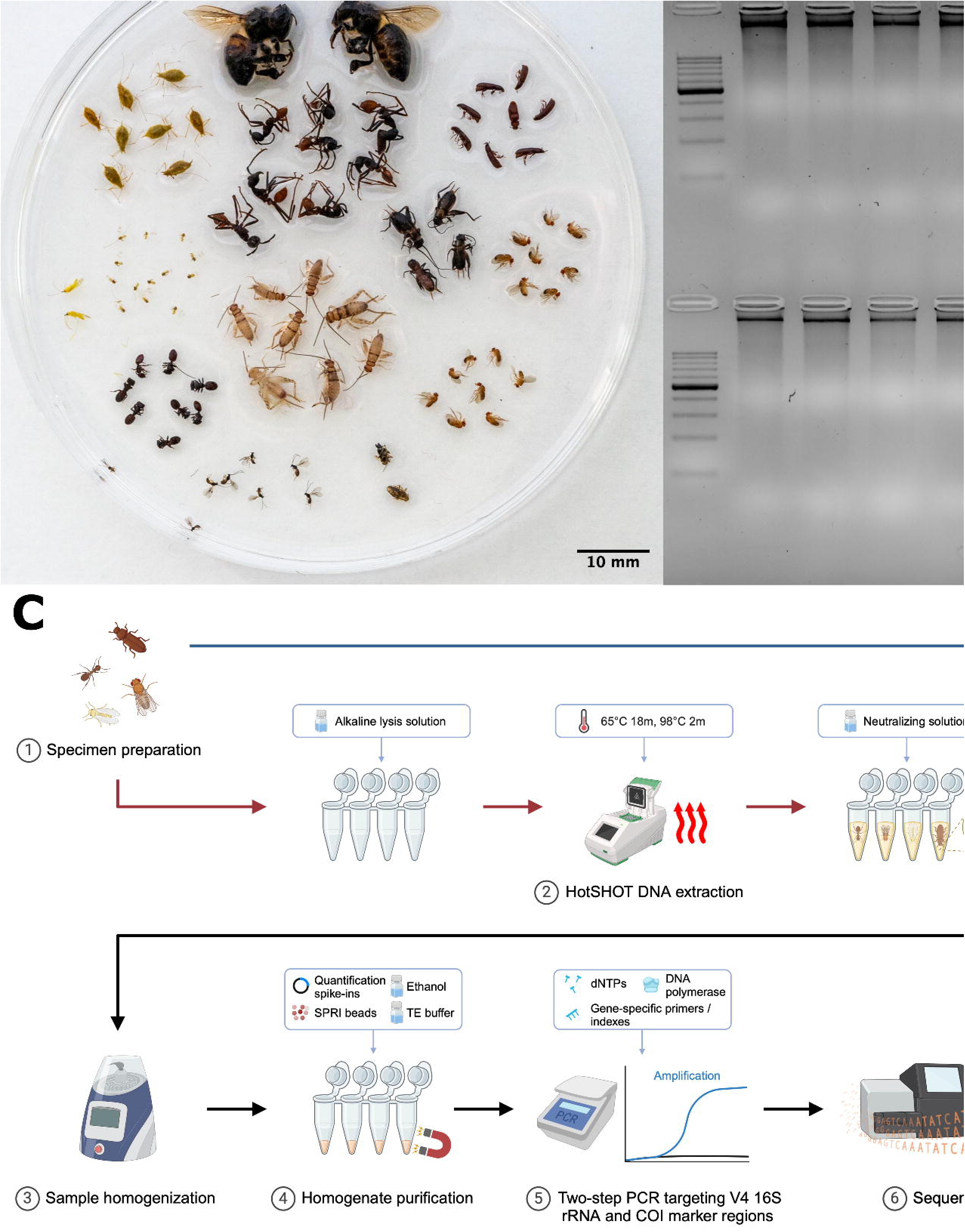
A. Representatives of the insect species used in the analysis. B. Comparison of DNA quality between samples treated with different extraction methods. C. Experimental workflow (Created with BioRender.com).

On the other hand, marker gene analysis may miss some taxa due to amplification biases and provide limited information on microbiome functional capabilities. Metagenomics, in contrast, can provide detailed information on the taxonomic position and function of more abundant symbionts, but its higher cost, computational demands, and analytical challenges limit broad implementation (Knight et al., 2018). For such methods, we want to select samples carefully. Generally, when addressing questions about microbiota in wild insects, we would want to combine methods that differ in resolution and per-sample cost by first surveying insect diversity broadly and then selecting specimens for a thorough characterization using more comprehensive methods. However, a key question is whether the different methods can plausibly be applied to the same specimen one after another.

In order to combine insect community characterization with microbiome characterization of the same insects, it is important to assess whether the voucher material generated during megabarcoding can be used for microbiome characterizing. Thus, we assessed the impact of the non-destructive DNA extraction method, HotSHOT, used in popular high-throughput barcoding protocols for insect diversity surveys (Srivathsan et al., 2021, 2023; Vasilita et al., 2024; Hu et al., 2024), on the reconstructed microbial community composition. Specifically, we asked whether microbial abundance estimates and the reconstructed bacterial community profile may change due to HotSHOT pre-treatment.

We did this by comparing quantitative V4 16S rRNA amplicon sequencing data for 286 specimens of 16 species, whether HotSHOT-treated or not (Fig. 1C). Our findings of the limited effects of HotSHOT/barcoding pre-treatment on insect microbial community profiles open up new avenues for monitoring the diversity and distribution of both insects and their microbial associates.

## Materials and methods

### Specimen collection and preparation

We selected 286 specimens belonging to 16 insect species representing six orders (Table 1). Species were selected based on the expectation of relative homogeneity in microbiota composition among individuals. They originated from standardized commercial or laboratory cultures, social insect colonies, clusters of hemipteran insects collected from the same plants and likely closely related or clonal, and in just one case (*D. hamata*), comprised different and likely unrelated individuals from a single population (Table 1). For three *Drosophila* species, we used distinct lines, one *Wolbachia*-positive and another *Wolbachia*-negative; for simplicity of the narrative, we will include these lines in the count of “species” further on. All the specimens of a species were at the same developmental stage but with some variation in body shape, cuticle properties, and size in at least some cases (Fig. 1A). The specimens were preserved in 95% ethanol and stored at -20L.

**Table 1.**
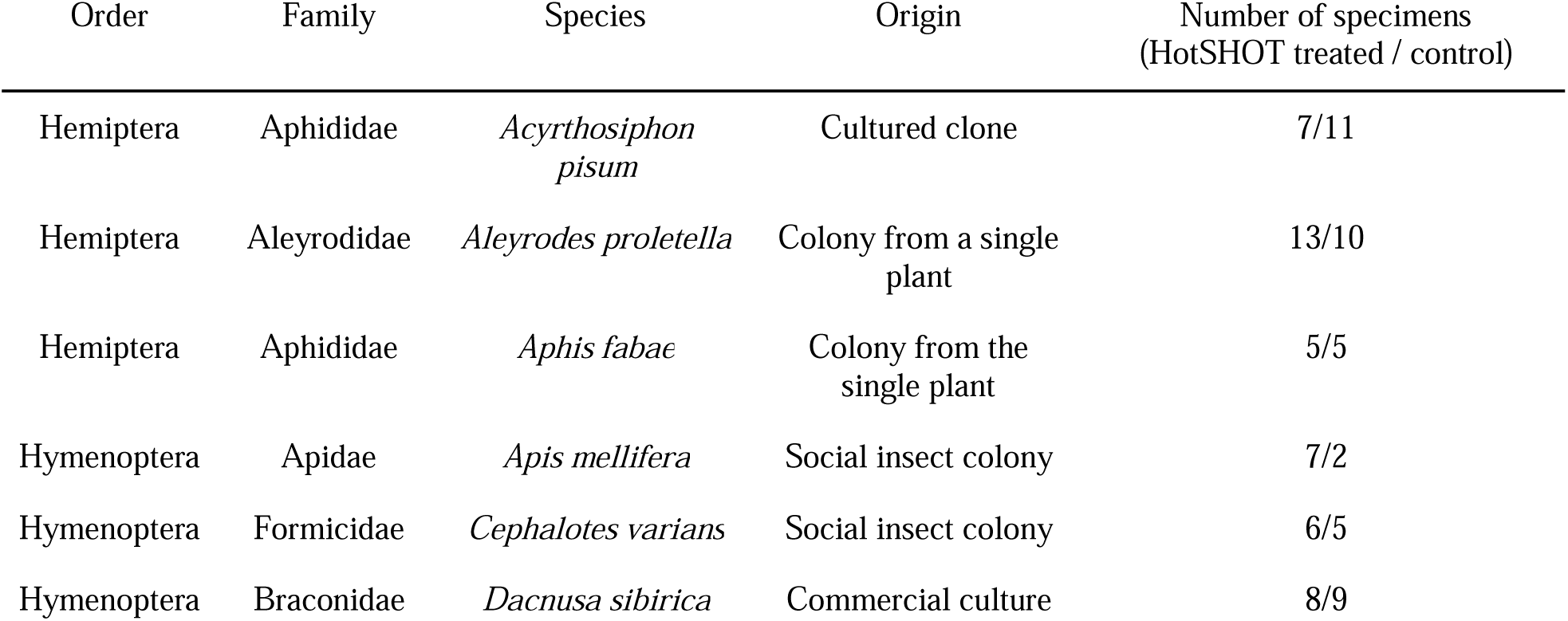
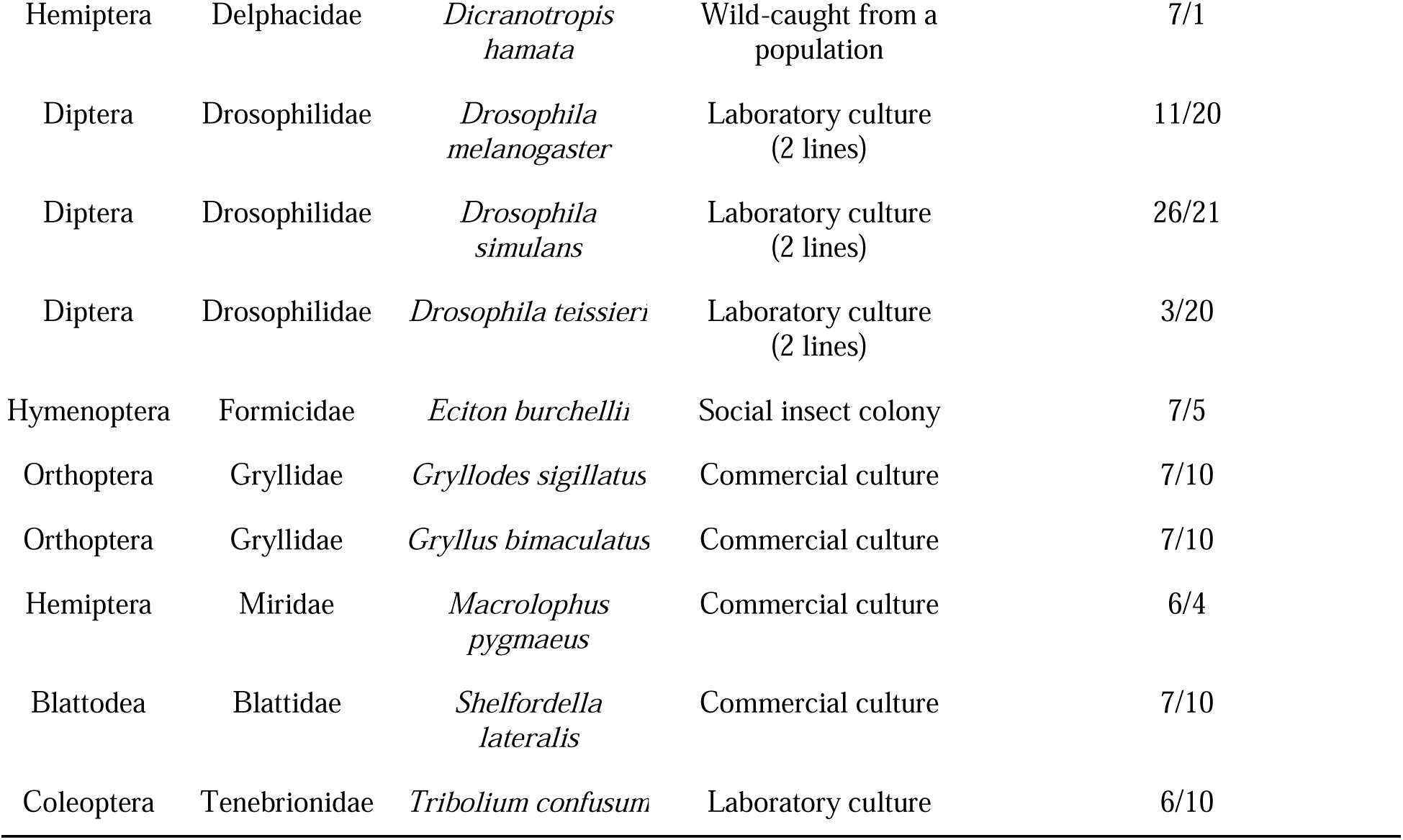
Insect species used in the study.

### Lysis and DNA extraction

Collected insects were divided into two groups, and from one group, DNA was extracted at the National University of Singapore using the HotSHOT method (Truett et al., 2000). The insects were placed in a 96-well plate filled with 10-15 µl of alkaline lysis solution (25 mM NaOH, 0.2 mM Na_2_EDTA, pH 12) individually and then incubated in a thermocycler for 20 minutes – 18 minutes at 65L and 2 minutes at 98L (Fig. 1C). In the case of larger insects, more lysis solution was added, although complete submersion was not necessary. Once heated, the DNA extract was neutralized by adding an equal volume of neutralization buffer (10-15 µl of 40 mM Tris-HCl) (Truett et al., 2000; Srivathsan et al., 2021). The DNA extract was then immediately used for the barcoding procedure, as described in Srivathsan et al., 2021, and the insects were shipped to Jagiellonian University (JU), Poland, for further processing.

From that point onwards, all insects (those HotSHOT-pretreated and untreated controls) underwent the same processing procedure. Before the procedure, the size of each individual was measured and categorized into five groups – 1 (up to 2 mm), 2 (up to 5 mm), 3 (up to 1 cm), 4 (up to 2 cm), 5 (more than 2 cm). For homogenization, the insects were placed in 2 ml tubes filled with 200 µl of a buffer mixture consisting of 195 µl ‘Vesterinen’ lysis buffer (0.4 M NaCl, 10 mM Tris-HCl, 2 mM EDTA pH 8, 2% SDS) (Aljanabi & Martinez, 1997; Vesterinen et al., 2016) and 5 µl proteinase K. More solutions were added if the sample was not completely submerged. After adding 2.8 and 0.5 mm ceramic beads, the tubes were grounded in the Omni Bead Raptor Elite homogenizer for two 30-second cycles, with the speed set to 5 m/s. Samples were then incubated in a thermal block at 55□ for 2 hours.

Once cooled, 40 µl of homogenate from each tube was transferred to a deep-well plate, where to each, we added a specific number of quantification spike-in (Table S1) – a linearized plasmid carrying an artificial 16S rRNA target Ec5001 (Tourlousse et al., 2017) suspended in 2 µl of TE buffer. The DNA was then purified with 80 µl of SPRI beads using a magnetic stand and washed twice with 80% ethanol. After dilution with 20.5 µl of TE buffer, 20 µl of the solution was transferred to a new 96-well plate, and the DNA concentration was measured with the Quant-iT PicoGreen kit.

### Library preparation and sequencing

The amplicon libraries were prepared following a custom two-step PCR protocol (Kolasa et al., 2023; Mulio et al., 2024). The first step involved simultaneous amplification of two marker regions: a V4 region of the 16S rRNA bacterial gene and a portion of an insect mitochondrial cytochrome oxidase I (COI) gene. We used template-specific primer pairs with Illumina adaptor tails: for the V4 region, 515F (GTGYCAGCMGCCGCGTAA) and 806R (GGACTACNVGTWTCTAAT) (Parada, Needham & Fuhrman, 2016), and for COI, BF3 (CCHGAYATRGCHTTYCCHCG) and BR2 (TCDGGRTGNCCRAARAAYCA) (Elbrecht et al., 2019). The PCR solution consisted of 5 µl of QIAGEN Multiplex Master Mix, a mix of primers at concentrations 2.5uM (COI) and 10uM (16S V4), 2 µl of DNA template, and 1 µl of water (final volume: 10 µl). The temperature program for the first round of PCR included the initial step of denaturation at 95°C for 15 minutes, followed by 25-27 cycles of denaturation (30s, 94°C), annealing (90s, 50°C) and extension (90s, 72°C) phases, and the final extension step (10m, 72°C). The products were checked on 2.5% agarose gel against positive and negative controls and cleaned with SPRI beads.

During the second indexing PCR, Illumina adapters and unique index pairs were added to the samples. The temperature program for this PCR remained the same, but the number of cycles was reduced to 7. As in previous steps, positive and negative controls were included to verify accuracy.

The libraries were pooled approximately equimolarly based on band intensity on agarose gels to ensure a roughly equal representation of each sample in the pool. After the last cleaning step with SPRI beads, the pools were ready for sequencing performed on an Illumina MiSeq v3 lane (2 × 300 bp reads) at the Institute of Environmental Sciences (JU).

### Bioinformatic processing

The bioinformatics analysis of the data was performed on a Unix cluster using a pipeline developed in the Symbiosis Evolution Research Group, combining custom Python scripts with already established bioinformatics tools. The pipeline documentation is available on the GitHub page (https://github.com/Symbiosis-JU/Bioinformatic-pipelines/blob/main/Example_analysis.md), and the versions of the scripts used in the current analysis are present in the GitHub repository – https://github.com/NAndriienko/HotSHOT-Microbiome-Project.

First, the amplicon data in FASTQ format were split into separate bins based on the primers using a dedicated script, which split the data based on used primer sequences into bins representing marker genes of interest (COI, 16SV4). As all of the specimens used in this study were of known species, we focused only on the 16S rRNA data in further steps in the analysis.

Initially, using PEAR, we assembled quality-filtered forward and reverse reads into contigs (Zhang et al., 2014). Next, contigs were de-replicated (Rognes et al., 2016) and denoised (Edgar, 2016) separately for every library to avoid losing information about rare genotypes that could happen during the denoising of the whole sequence set at once (Prodan et al., 2020). The sequences were then screened for chimeras using USEARCH and classified by taxonomy using the SINTAX algorithm and customized SILVA database (version 138 SSU) (Quast et al., 2013). Finally, the sequences were clustered at a 97% identity level using the UPARSE-OTU algorithm implemented in USEARCH. The tables with two levels of classification were produced: ASVs (Amplicon Sequencing Variant) (also known as zOTUs - zero-radius Operational Taxonomic Units) describing genotypic diversity and OTUs (Operational Taxonomic Units) – clustering genotypes based on a similarity threshold.

Bacterial 16S rRNA gene data were screened for putative DNA extraction and PCR reagent contaminants using negative controls (blanks) for each laboratory step as a reference. We first used taxonomy classification information to remove genotypes classified as chloroplasts, mitochondria, Archea, or chimers. Next, we calculated relative abundances and used ratios of each genotype presented in blank and experimental libraries to accurately assign genotypes as putative actual insect-associated microbes or PCR or extraction contaminants (Table S2).

Additionally, reads identified as quantitative spike-ins were used to reconstruct bacterial absolute abundances in the processed insects. Specifically, the symbiont-to-extraction spike-in ratio, multiplied by the number of extraction spike-in copies and the proportion of the homogenate, allowed us to estimate amplifiable bacterial 16S rRNA copy numbers in the homogenized specimens.

Finally, manual analysis was conducted to remove controls, samples with incorrect indexes or zero abundance of bacteria and create the dataset used in the statistical analysis.

### Statistical analysis and visualization

Statistical analysis was performed using RStudio version 2023.03.1+446 (R Core Team, 2023) and QIIME2 2023.2 (Bolyen et al., 2019) software. Inkscape 1.2.2 (Inkscape Project, 2022) was used to modify generated plots and visualizations.

One-way ANOVA with random effects on insect species was used for the absolute abundance comparison between groups treated with HotSHOT and those that were only homogenized. The base and ‘nlme’ (Pinheiro, Bates & R Core Team, 2023) packages were utilized for this step. The analysis was visualized with the usage of ‘ggplot2’ (Wickham, 2016), ‘dplyr’ (Wickham et al., 2023), ‘RColorBrewer’ (Neuwirth, 2022) and ‘phyloseq’ (McMurdie & Holmes, 2013) packages. A biodiversity assessment was performed in QIIME2 on the Unix cluster. Firstly, the decontaminated OTU table, bacteria taxonomy, and file with bacterial OTU sequences in fasta format were imported into QIIME2 as artifacts to allow further transformation and tracking of the origin of the output files of the analysis (McDonald et al., 2012). Next, the obtained feature table was filtered separately for each insect species so the differences would not disturb the analysis.

As diversity metrics are sensitive to different sampling depths in the groups, we standardized the samples by rarefying them (Weiss et al., 2017). Rarefaction level was chosen individually based on the generated summary for these tables, balancing between retaining the highest amount of samples and the percentage of the features left for analysis.

The alpha- and beta-diversity indexes, calculated using the q2-diversity plugin, were used to compare the microbiome composition between the methods for each species. Alpha diversity refers to the diversity within the samples (Whittaker, 1972), and Shannon’s entropy (Shannon, 1948), which combines richness and evenness evaluation in a single metric and provides a comprehensive assessment of diversity, was chosen for its assessment. The calculated indexes were compared between the groups using the Kruskal-Wallis statistical test (Kruskal & Wallis, 1952).

The Bray-Curtis dissimilarity (Sørensen, 1948) index was chosen for beta-diversity analysis. This index calculates the dissimilarity between communities based on the presence and abundance of different features. It considers both the presence/absence and the relative abundances of features, providing a quantitative measure of compositional differences between samples.

To visualize the results and explore the beta-diversity patterns, the Principle Coordinate Analysis (PCoA) (Halko et al., 2011; Legendre & Legendre, 2012) was performed using the Emperor tool (Vázquez-Baeza et al., 2013). Moreover, the PERMANOVA and PERMDISP tests were conducted to analyze the statistical trends (Anderson, 2001).

## Results

We obtained reliable data for a total of 286 biological samples. We analyzed and interpreted them using an additional 16 negative control samples of different types. Each sample yielded a minimum of 55 16S rRNA reads, ranging from 55 to 93 394 reads, and a mean of 24 100 reads (Table S1).

### The effects of treatment on the microbiome absolute abundance

The quantification spikein was present in every amplicon library. In experimental samples, the ratio of bacterial reads not classified as contamination to Ec5001 (extraction plasmid) reads ranged from 0.0114 (in *Dacnusa sibirica*) to 16187.5 (in *Drosophila simulans*). With 10,000 copies of artificial amplification target Ec5001 added to 20% of insect homogenate (in most cases), and assuming no amplification bias among different types of templates, this translates to between 697 and 2,062,562,500 copies of bacterial 16S rRNA per insect. When comparing estimated microbial abundances across species and treatments (two-way ANOVA using log-transformed data), we found significant differences between treatments (F = 159.86, p < 0.0001). However, the effect of HotSHOT treatment varied among the species (Fig. 2). We tested the treatment effect in each of the 16 species.

**Figure 2.**
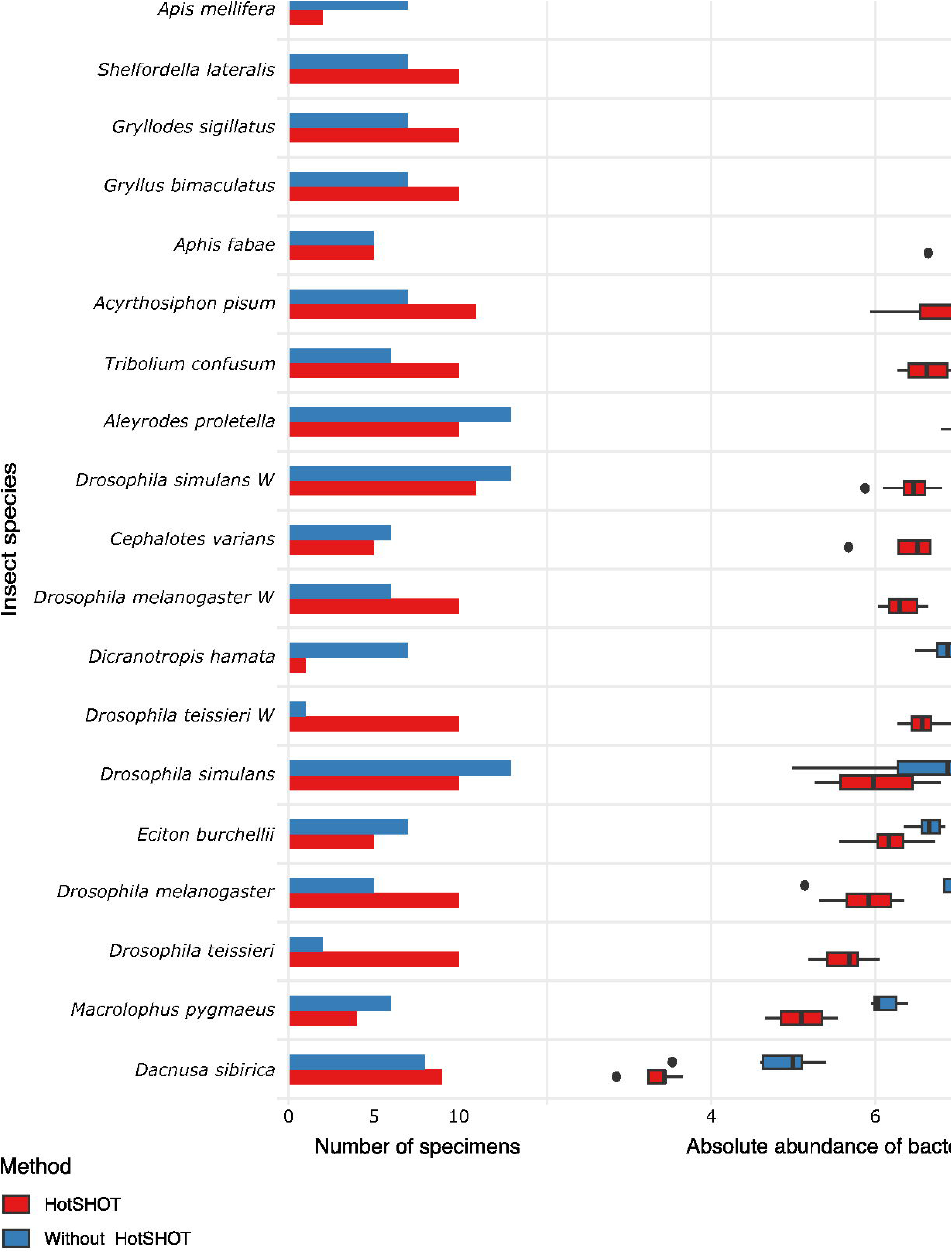
Comparison of the absolute abundance of bacteria between specimens treated with HotSHOT and control specimens among insect species.

Further analysis using the Kruskal-Wallis test showed that bacterial abundance was significantly lower after HotSHOT in 13 out of 19 species (Suppl. Table S3). We also checked if the treatment effect was connected with the insects’ size, and test results indicate that it was similarly significant for all size categories (Suppl. Table 3). On average, the estimated bacterial absolute abundance decreased 15-fold following HotSHOT. We conclude that these values represent the actual decrease in the concentration of amplifiable gene targets during incubation in the hot alkaline buffer.

### Microbiome diversity analysis

We obtained biologically realistic data for all species (Fig. 3, Suppl. Tables S4-5). As expected, we observed *Acetobacter* and *Lactobacillus* as the dominant symbionts of cultured *Drosophila* species (Wong, Ng & Douglas, 2011), *Buchnera* as the dominant symbiont in two aphids (Douglas, 2003), and well-known members of gut microbiota of honeybees (Engel & Moran, 2013) and *Cephalotes* (Hu et al., 2018) and *Eciton* ants (Mendoza-Guido et al., 2023). Facultative endosymbiont *Wolbachia* was present in 5 species and often dominated their microbial communities, whereas *Cardinium* dominated the microbiota of *D. hamata* planthopper. In cultured *Gryllodes sigillatus* and *Gryllus bimaculatus*, we found relatively complex microbiota comprising relatively widespread bacterial genera as well as some specialists (*Blattabacterium* in *S. lateralis*) at variable abundances. In the species with the least abundant microbiota, parasitic wasp *D. sibirica*, microbiota composition was highly variable.

**Figure 3.**
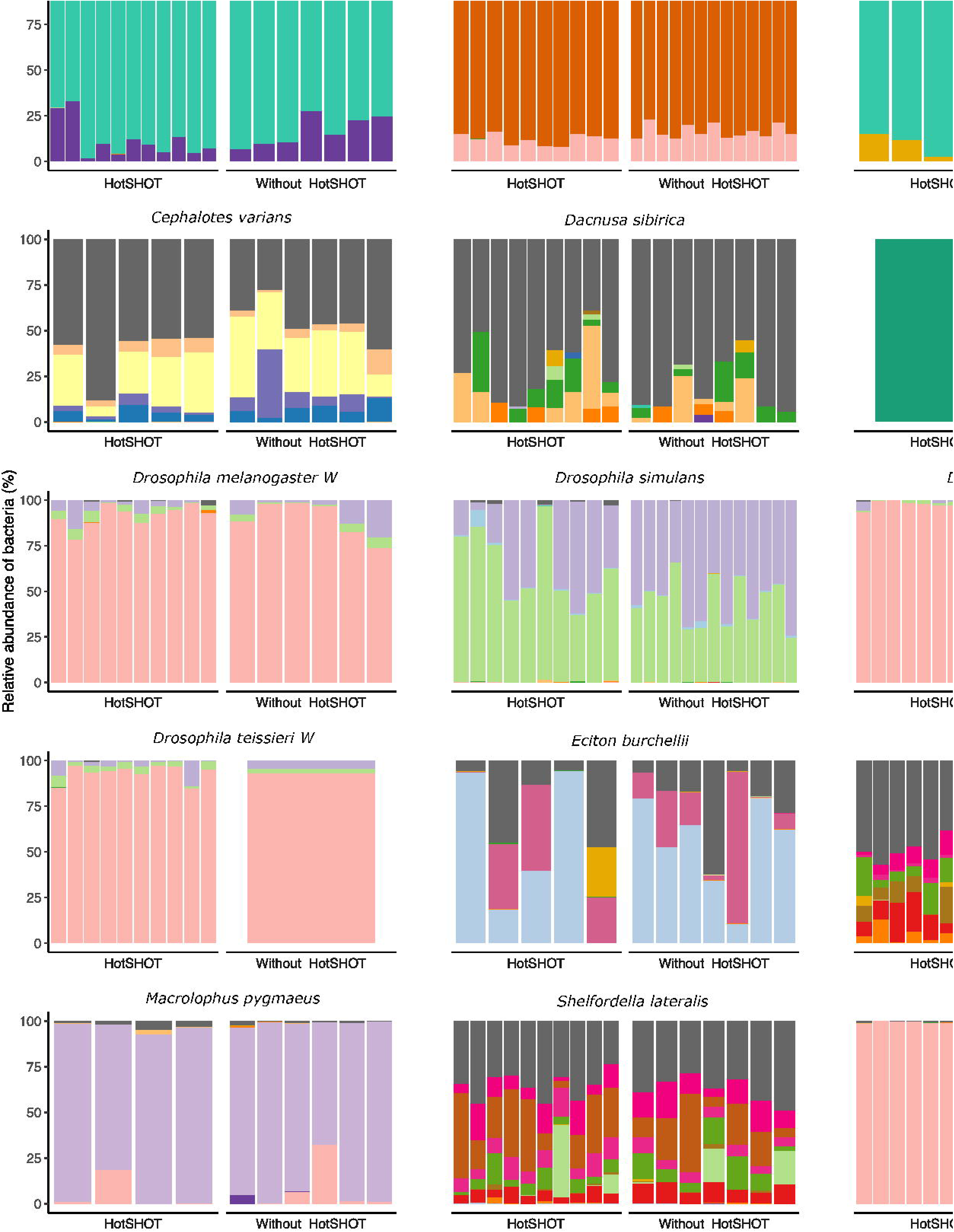
Relative abundances of bacterial genera in 19 lines of 16 insect species. Bars represent individual insects, whether HotSHOT-treated or control, of that line. Colors represent microbial genera above 5%, and gray represents the summed abundance of all other genera combined.

The differences in microbiota profiles for individuals representing the same lines or species but exposed to different treatments are not obvious at first sight. In most cases, we observed no such effect when conducting more formal analyses. We observed differences in richness and evenness in the distribution of features in three species (*Aleyrodes proletella*, *Gryllodes sigillatus,* and *Tribolium confusum*), which were expressed through significant changes in Shannons’ entropy values (Suppl. Table S6). Beta diversity analysis, conducted using the Bray-Curtis index, revealed significant differences in microbial community composition in 4 other species (*Aleyrodes proletella*, *Cephalotes varians*, *Dacnusa sibirica*, and *Drosophila teissieri*). These differences in the presence and relative abundance of features can be observed on PCoA plots for these species (Suppl. Fig. S1), where the first axis explains from 15.47% to 86.16% of diversity.

The comparison of average relative abundances of dominant bacterial clades (Fig. 4) shows that the bacterial taxa mostly overlap between the different methods, despite fluctuations in the proportions of some bacterial taxa, but they are not constant. We observed minor changes in the proportions of primary insect symbionts. However, the effects of pre-barcoding are not entirely clear as there are also cases of increasing proportions, and all of these bacteria were present in trace amounts in the samples.

**Figure 4.**
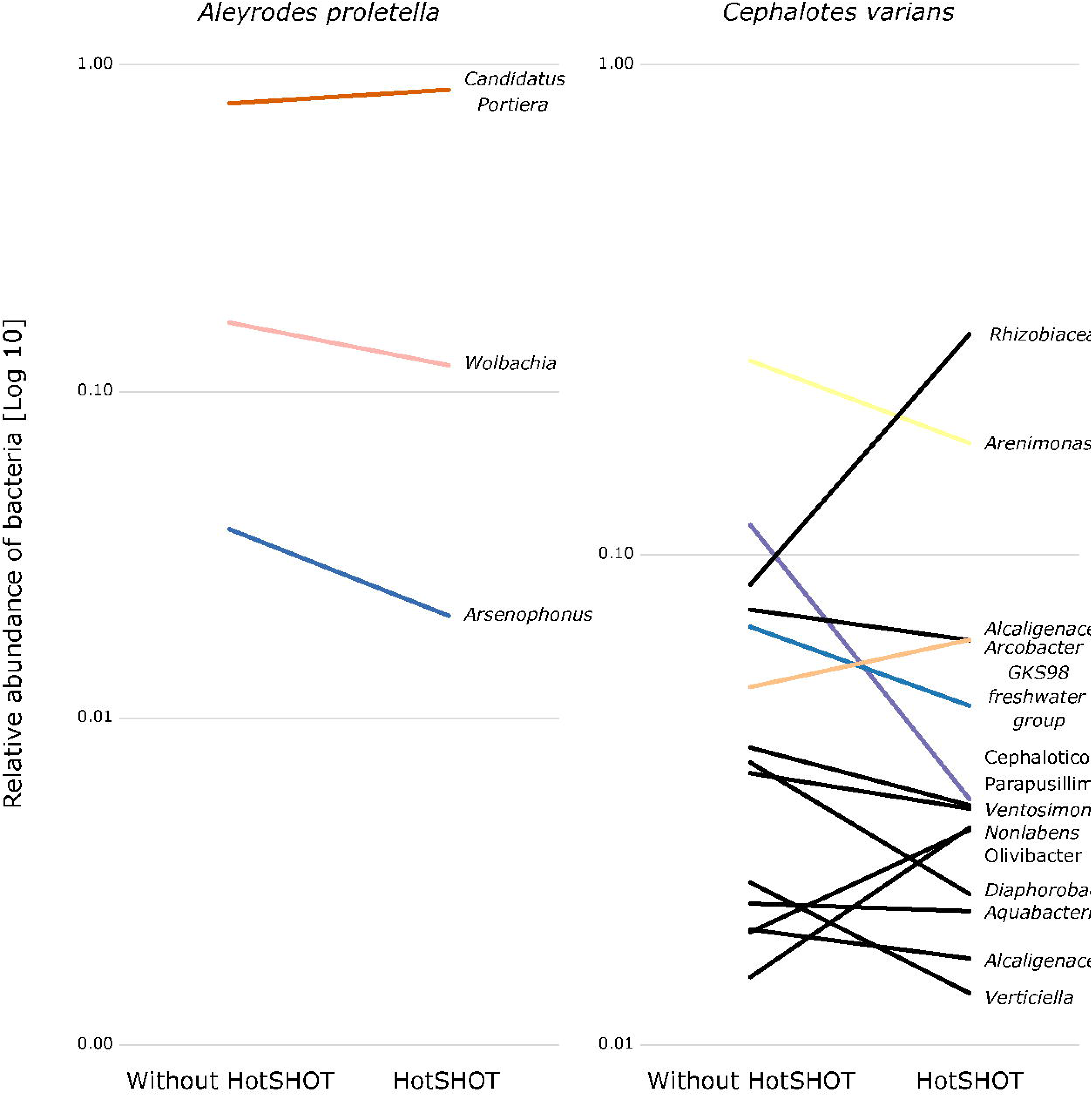
Changes in the relative representation of microbial genera above 1% among treatments.

Overall, the microbial communities were dominated by known bacterial commensals of insects and, in many cases, by commensals characteristic of the particular species. In general, dominant bacteria were present in all individuals of a species, although their relative abundance varied slightly, which might reflect natural variation between specimens. Significant differences in composition were not systematic and were related to changes in rarely present features rather than changes in major symbionts. Thus, no overall difference in composition between pre-barcoded and control samples was observed.

## Discussion

The DNA extraction that involves specimen incubation in the hot alkaline buffer, called HotSHOT, affects the DNA integrity and, consequently, the microbial DNA yield within the treated specimens. This results in decreased absolute abundance estimates of 16S rRNA copies in processed specimens, which is presumably a direct consequence of DNA degradation caused by the HotSHOT treatment. Nevertheless, the reconstructed microbial community composition, especially the reconstructed presence of abundant microbial clades, which are likely to play the most significant roles in insect biology, was not substantially affected. Thus, even though the barcoding process disrupts our ability to estimate **the amounts** of bacteria colonizing insect bodies, it does not seem to substantially bias conclusions about the **identity** of these bacteria.

### Effects on microbial abundance

Insects and other organisms differ dramatically in the abundance of microorganisms they host, and these differences often correlate with the microbes’ importance in insect biology (Hammer, Sanders & Fierer, 2019). Abundance is a critical aspect of host-microbiome relationships. Assuming all else equal, the greater number of microbial cells should translate to their stronger effect, driven by higher nutritional demands and greater amounts of biologically active compounds produced. However, the effects may not be linear due to the bacterial ability to detect their abundance and alter biological activity in response (*quorum sensing*) (Miller & Bassler, 2001). For example, some microbial pathogens may delay the production of virulence factors until they sense abundance sufficient for overwhelming host defenses (Munoz et al., 2020).

On the other hand, in *Sodalis praecaptivus* – a versatile opportunist that seems representative of the ancestral state of many heritable symbionts, *quorum sensing* attenuates virulence, facilitating a long-lasting and benign association (Enomoto et al., 2017). Further, one could imagine that symbionts with significant biological effects would be promoted either by host- or microbe-encoded mechanisms and reach higher abundance. From a more technical perspective, abundance also determines the precision of microbiome composition reconstruction. We know that low-bacterial-abundance samples are much more prone to contamination from reagents and other sources (Salter et al., 2014), likely leading to erroneous conclusions. We can be much more confident about the presence of abundant microbes.

For these reasons, it is important to estimate the absolute abundance of the microbiome rather than just report the relative abundance of microbial clades. Unfortunately, with some notable exceptions (Hammer et al., 2017; Sanders et al., 2017; Ravenscraft et al., 2019; Surmacz et al., 2024), it is rarely done in experimental studies. We argue that the available tools for abundance estimation, including quantitative PCRs and spike-ins (Props et al., 2017; Tourlousse et al., 2017; Jian et al., 2020; Harrison et al., 2021), should become a part of standard microbiome analyses. At the same time, we must be aware that aggressive treatments, such as incubation in an alkaline buffer at high temperatures, may lead to the degradation of the substantial share of the available DNA and disrupt or complicate abundance reconstruction. Changes to the HotSHOT protocol, such as reduced incubation time, will likely limit DNA degradation while yielding sufficient amounts for barcoding analyses. Indeed, we have recently reduced the routine incubation time to 5 minutes without compromising barcoding results (V. Andriienko & A. Michalik, unpublished).

### Effects on microbial composition

Organisms differ in which microorganisms they host, and different microbial functional categories, clades, and even strains have very different effects on host biology and evolution (Bourtzis & Miller, 2003).

Our study used insect species hosting a broad range of microbial functional categories and taxonomic clades, reflecting the broad range of host ecological niches. Our results are congruent with *a priori* expectations about symbionts present in these species, and HotSHOT treatment has not affected the detection of their dominant microbes. We conclude that our ability to detect abundant bacteria, including nutritional and facultative endosymbionts and specialized gut bacteria known to have major effects on the host ecology, is not being altered by HotSHOT pre-treatment. On the other hand, the ability to detect less abundant microbes may be reduced. However, given the known issues with reagents- and cross-contamination, the reliability of their detection may be limited regardless, even when appropriate controls are used. Such low-abundance microbes, even if reliably detected, are also less likely to form biologically relevant associations significant to the host.

Statistical analyses indicate that in a few cases, the HotSHOT treatment elevates the abundance of some bacteria taxa. We interpret those results as an outcome of the reduction of symbiont DNA amounts caused by the HotSHOT treatment, with more pronounced contamination. However, it is on a relatively low level and does not change the overall composition of the insect microbiome. Nevertheless, this fact should not be trivialized, and appropriate negative controls should be implemented and processed to filter out introduced bacterial signal effectively.

Considering this, the HotSHOT treatment may confound some of the conventional approaches to microbiome analysis, including altering zOTU or OTU counts, rarefaction, or diversity index comparisons. While these approaches may be considered the “gold standard” nowadays, we argue that researchers must carefully consider the biological value and relevance of the information they provide for their system, taking into consideration reagent contamination and other challenges. This is particularly relevant for small organisms such as insects, with often low overall bacterial abundance or dominated by one of few abundant microbes and with the remainder at low abundance.

While this should be decided on a case-by-case basis, we think that analyses focused on abundant insect symbionts – unaffected by the HotSHOT treatment – provide much more biologically relevant information.

### Broader context

The methods for biodiversity discovery have been shifting, with approaches such as ultra-throughput individual barcoding, or alternatively, metabarcoding, rapidly gaining in popularity. International projects such as BOLD are already processing millions of insect specimens to discover global insect diversity. Thus, we have a growing amount of individuals processed with non-destructive DNA extraction techniques. There is an assumption that these insects remain suitable for further morphological or DNA-based characterization. However, there are limited examples of successful usage of such pre-processed specimens for addressing further questions. Our work demonstrates clearly that the associations with some of the most important players in insect biology – symbiotic microorganisms – can be reliably reconstructed from such material. This opens up exciting avenues for microbiota study across large numbers of pre-barcoded wild-caught specimens and cost-effective reconstruction of broad microbiome-related patterns. With bacterial symbionts increasingly regarded as important players in insect biology and a potential source of rapid insect adaptation to changing environments, we advocate that microbiome screening should become a standard procedure in insect biodiversity studies (Łukasik & Kolasa, 2023). Hence, rather than attempting labor- and cost-intensive additional sampling for fresh material, relying on microbiome typing on pre-barcoded material could be an extremely frugal way of conducting science. Simultaneously, our approach confirms that the material will likely be suitable for other DNA-based approaches aiming to unravel mechanisms that shape insect diversity and biology.

## Conclusions

Biodiversity and microbiome researchers have multiple tools that vary in their information output, per-sample cost, and plausible throughput. When addressing biological questions, it is essential to balance these criteria. However, our results suggest that we do not need to limit ourselves to just one tool. On the contrary, by serially applying different methods to the same specimens, we can combine the breadth of extremely cost-effective approaches, such as barcoding, with much deeper insights that could be obtained from multi-gene amplicon sequencing and, likely, also genomics tools. Combining these different tools could become a new paradigm for biodiversity studies in the turbulent era of the Anthropocene.

## Data availability

All data underlying this study have been deposited in the Sequence Read Archive (SRA) of the National Center for Biotechnology Information (NCBI) under BioProject Accession no. PRJNA1102268

## Acknowledgments

We thank Brandon Cooper and Jacob Russell for providing some of the specimens that were used in this study. The project was supported by the Polish National Agency for Academic Exchange grant PPN/PPO/2018/1/00015, the Polish National Science Centre grants 2018/31/B/NZ8/01158 and 2021/43/B/NZ8/03376, the National Institute of General Medical Sciences of the NIH award number R35GM124701 to Brandon S. Cooper (insect culture), and Jagiellonian University POB BioS minigrant (ID: B.1.11.2020.101).

## References

Aljanabi SM, Martinez I. 1997. Universal and rapid salt-extraction of high quality genomic DNA for PCR-based techniques. Nucleic Acids Research 25:4692–4693. DOI: 10.1093/nar/25.22.4692.

Anderson MJ. 2001. A new method for non-parametric multivariate analysis of variance. Austral Ecology 26:32–46. DOI: 10.1111/j.1442-9993.2001.01070.pp.x.

Bolyen E, Rideout JR, Dillon MR, Bokulich NA, Abnet CC, Al-Ghalith GA, Alexander H, Alm EJ, Arumugam M, Asnicar F, Bai Y, Bisanz JE, Bittinger K, Brejnrod A, Brislawn CJ, Brown CT, Callahan BJ, Caraballo-Rodríguez AM, Chase J, Cope EK, Da Silva R, Diener C, Dorrestein PC, Douglas GM, Durall DM, Duvallet C, Edwardson CF, Ernst M, Estaki M, Fouquier J, Gauglitz JM, Gibbons SM, Gibson DL, Gonzalez A, Gorlick K, Guo J, Hillmann B, Holmes S, Holste H, Huttenhower C, Huttley GA, Janssen S, Jarmusch AK, Jiang L, Kaehler BD, Kang KB, Keefe CR, Keim P, Kelley ST, Knights D, Koester I, Kosciolek T, Kreps J, Langille MGI, Lee J, Ley R, Liu Y-X, Loftfield E, Lozupone C, Maher M, Marotz C, Martin BD, McDonald D, McIver LJ, Melnik AV, Metcalf JL, Morgan SC, Morton JT, Naimey AT, Navas-Molina JA, Nothias LF, Orchanian SB, Pearson T, Peoples SL, Petras D, Preuss ML, Pruesse E, Rasmussen LB, Rivers A, Robeson MS, Rosenthal P, Segata N, Shaffer M, Shiffer A, Sinha R, Song SJ, Spear JR, Swafford AD, Thompson LR, Torres PJ, Trinh P, Tripathi A, Turnbaugh PJ, Ul-Hasan S, vander Hooft JJJ, Vargas F, Vázquez-Baeza Y, Vogtmann E, von Hippel M, Walters W, Wan Y, Wang M, Warren J, Weber KC, Williamson CHD, Willis AD, Xu ZZ, Zaneveld JR, Zhang Y, Zhu Q, Knight R, Caporaso JG. 2019. Reproducible, interactive, scalable and extensible microbiome data science using QIIME 2. Nature biotechnology 37:852–857. DOI: 10.1038/s41587-019-0209-9.

Bourtzis K, Miller TA. 2003. Insect Symbiosis. Boca Raton: CRC Press. DOI: 10.1201/9780203009918.

Buchner D, Sinclair JS, Ayasse M, Beermann A, Buse J, Dziock F, Enss J, Frenzel M, Hörren T, Li Y, Monaghan MT, Morkel C, Müller J, Pauls SU, Richter R, Scharnweber T, Sorg M, Stoll S, Twietmeyer S, Weisser WW, Wiggering B, Wilmking M, Zotz G, Gessner MO, Haase P, Leese F. 2024. Upscaling biodiversity monitoring: Metabarcoding estimates 31,846 insect species from Malaise traps across Germany. 2023.05.04.539402. DOI: 10.1101/2023.05.04.539402.

Chua PYS, Bourlat SJ, Ferguson C, Korlevic P, Zhao L, Ekrem T, Meier R, Lawniczak MKN. 2023. Future of DNA-based insect monitoring. Trends in Genetics 39:531–544. DOI: 10.1016/j.tig.2023.02.012.

Douglas A. 2003. *Buchnera* bacteria and other symbionts of aphids. In: Insect symbiosis.

Dwyer G, Dushoff J, Yee SH. 2004. The combined effects of pathogens and predators on insect outbreaks. Nature 430:341–345. DOI: 10.1038/nature02569.

Edgar RC. 2016. UNOISE2: improved error-correction for Illumina 16S and ITS amplicon sequencing. :081257. DOI: 10.1101/081257.

Elbrecht V, Braukmann TWA, Ivanova NV, Prosser SWJ, Hajibabaei M, Wright M, Zakharov EV, Hebert PDN, Steinke D. 2019. Validation of COI metabarcoding primers for terrestrial arthropods. PeerJ 7:e7745. DOI: 10.7717/peerj.7745.

Engel P, Moran NA. 2013. Functional and evolutionary insights into the simple yet specific gut microbiota of the honey bee from metagenomic analysis. Gut Microbes 4:60–65. DOI: 10.4161/gmic.22517.

Enomoto S, Chari A, Clayton AL, Dale C. 2017. Quorum sensing attenuates virulence in *Sodalis praecaptivus*. Cell Host & Microbe 21:629–636.e5. DOI: 10.1016/j.chom.2017.04.003.

Ferrari J, Vavre F. 2011. Bacterial symbionts in insects or the story of communities affecting communities. Philosophical Transactions of the Royal Society B: Biological Sciences 366:1389–1400. DOI: 10.1098/rstb.2010.0226.

Geiger M, Moriniere J, Hausmann A, Haszprunar G, Wägele W, Hebert P, Rulik B. 2016. Testing the Global Malaise Trap Program – How well does the current barcode reference library identify flying insects in Germany? Biodiversity Data Journal 4:e10671. DOI: 10.3897/BDJ.4.e10671.

Gibson J, Shokralla S, Porter TM, King I, van Konynenburg S, Janzen DH, Hallwachs W, Hajibabaei M. 2014. Simultaneous assessment of the macrobiome and microbiome in a bulk sample of tropical arthropods through DNA metasystematics. Proceedings of the National Academy of Sciences 111:8007–8012. DOI: 10.1073/pnas.1406468111.

Halko N, Martinsson P-G, Shkolnisky Y, Tygert M. 2011. An Algorithm for the Principal Component Analysis of Large Data Sets. SIAM Journal on Scientific Computing. DOI: 10.1137/100804139.

Hammer TJ, Janzen DH, Hallwachs W, Jaffe SP, Fierer N. 2017. Caterpillars lack a resident gut microbiome. Proceedings of the National Academy of Sciences of the United States of America 114:9641–9646. DOI: 10.1073/pnas.1707186114.

Hammer TJ, Sanders JG, Fierer N. 2019. Not all animals need a microbiome. FEMS Microbiology Letters 366:fnz117. DOI: 10.1093/femsle/fnz117.

Harrison JG, John Calder W, Shuman B, Alex Buerkle C. 2021. The quest for absolute abundance: The use of internal standards for DNA-based community ecology. Molecular Ecology Resources 21:30–43. DOI: 10.1111/1755-0998.13247.

Hu F-S, Arriaga-Varela E, Biffi G, Bocák L, Bulirsch P, Damaška AF, Frisch J, Hájek J, Hlaváč P, Ho B-H, Ho Y-H, Hsiao Y, Jelínek J, Klimaszewski J, Kundrata R, Löbl I, Makranczy G, Matsumoto K, Phang G-J, Ruzzier E, Schülke M, Švec Z, Telnov D, Tseng W-Z, Yeh L-W, Le M-H, Fikáček M. 2024. Forest leaf litter beetles of Taiwan: first DNA barcodes and first insight into the fauna. Deutsche Entomologische Zeitschrift 71:17–47. DOI: 10.3897/dez.71.112278.

Hu Y, Sanders JG, Łukasik P, D’Amelio CL, Millar JS, Vann DR, Lan Y, Newton JA, Schotanus M, Kronauer DJC, Pierce NE, Moreau CS, Wertz JT, Engel P, Russell JA. 2018. Herbivorous turtle ants obtain essential nutrients from a conserved nitrogen-recycling gut microbiome. Nature Communications 9:964. DOI: 10.1038/s41467-018-03357-y.

Husnik F. 2018. Host–symbiont–pathogen interactions in blood-feeding parasites: nutrition, immune cross-talk and gene exchange. Parasitology 145:1294–1303. DOI: 10.1017/S0031182018000574.

Inkscape Project. 2022. Inkscape.

Iwaszkiewicz-Eggebrecht E, Granqvist E, Buczek M, Prus M, Kudlicka J, Roslin T, Tack AJM, Andersson AF, Miraldo A, Ronquist F, Łukasik P. 2023. Optimizing insect metabarcoding using replicated mock communities. Methods in Ecology and Evolution 14:1130–1146. DOI: 10.1111/2041-210X.14073.

Jian C, Luukkonen P, Yki-Järvinen H, Salonen A, Korpela K. 2020. Quantitative PCR provides a simple and accessible method for quantitative microbiota profiling. PLOS ONE 15:e0227285. DOI: 10.1371/journal.pone.0227285.

Knight R, Vrbanac A, Taylor BC, Aksenov A, Callewaert C, Debelius J, Gonzalez A, Kosciolek T, McCall L-I, McDonald D, Melnik AV, Morton JT, Navas J, Quinn RA, Sanders JG, Swafford AD, Thompson LR, Tripathi A, Xu ZZ, Zaneveld JR, Zhu Q, Caporaso JG, Dorrestein PC. 2018. Best practices for analysing microbiomes. Nature Reviews Microbiology 16:410–422. DOI: 10.1038/s41579-018-0029-9.

Kolasa M, Kajtoch Ł, Michalik A, Maryańska-Nadachowska A, Łukasik P. 2023. Till evolution do us part: The diversity of symbiotic associations across populations of *Philaenus* spittlebugs. Environmental Microbiology 25:2431–2446. DOI: 10.1111/1462-2920.16473.

Kolasa M, Ścibior R, Mazur MA, Kubisz D, Dudek K, Kajtoch Ł. 2019. How Hosts Taxonomy, Trophy, and Endosymbionts Shape Microbiome Diversity in Beetles. Microbial Ecology 78:995–1013. DOI: 10.1007/s00248-019-01358-y.

Kruskal WH, Wallis WA. 1952. Use of ranks in one-criterion variance analysis. Journal of the American statistical Association 47:583–621. DOI: 10.1080/01621459.1952.10483441.

Langille MGI, Zaneveld J, Caporaso JG, McDonald D, Knights D, Reyes JA, Clemente JC, Burkepile DE, Vega Thurber RL, Knight R, Beiko RG, Huttenhower C. 2013. Predictive functional profiling of microbial communities using 16S rRNA marker gene sequences. Nature Biotechnology 31:814–821. DOI: 10.1038/nbt.2676.

Legendre P, Legendre L. 2012. Numerical Ecology. In: Elsevier, 499.

Lemoine MM, Engl T, Kaltenpoth M. 2020. Microbial symbionts expanding or constraining abiotic niche space in insects. Current Opinion in Insect Science 39:14–20. DOI: 10.1016/j.cois.2020.01.003.

Losey JE, Vaughan M. 2006. The Economic Value of Ecological Services Provided by Insects. BioScience 56:311–323. DOI: 10.1641/0006-3568(2006)56[311:TEVOES]2.0.CO;2.

Łukasik P, Kolasa MR. 2023. With a little help from my friends: the roles of microbial symbionts in insect populations and communities.

McDonald D, Clemente JC, Kuczynski J, Rideout JR, Stombaugh J, Wendel D, Wilke A, Huse S, Hufnagle J, Meyer F, Knight R, Caporaso JG. 2012. The Biological Observation Matrix (BIOM) format or: how I learned to stop worrying and love the ome-ome. GigaScience 1:2047–217X-1–7. DOI: 10.1186/2047-217X-1-7.

McFall-Ngai M, Hadfield MG, Bosch TCG, Carey HV, Domazet-Lošo T, Douglas AE, Dubilier N, Eberl G, Fukami T, Gilbert SF, Hentschel U, King N, Kjelleberg S, Knoll AH, Kremer N, Mazmanian SK, Metcalf JL, Nealson K, Pierce NE, Rawls JF, Reid A, Ruby EG, Rumpho M, Sanders JG, Tautz D, Wernegreen JJ. 2013. Animals in a bacterial world, a new imperative for the life sciences. Proceedings of the National Academy of Sciences 110:3229–3236. DOI: 10.1073/pnas.1218525110.

McMurdie PJ, Holmes S. 2013. phyloseq: An R Package for Reproducible Interactive Analysis and Graphics of Microbiome Census Data. PLoS ONE 8:e61217. DOI: 10.1371/journal.pone.0061217.

Meier R, Wong W, Srivathsan A, Foo M. 2016. $1 DNA barcodes for reconstructing complex phenomes and finding rare species in specimen-rich samples. Cladistics 32:100–110. DOI: 10.1111/cla.12115.

Mendoza-Guido B, Rodríguez-Hernández N, Ivens ABF, Beeren C von, Murillo-Cruz C, Zuniga-Chaves I, Łukasik P, Sanchez E, Kronauer DJC, Pinto-Tomás AA. 2023. Low diversity and host specificity in the gut microbiome community of *Eciton* army ants (Hymenoptera: Formicidae: Dorylinae) in a Costa Rican rainforest. Myrmecological News 33:19–34. DOI: 10.25849/myrmecol.news_033:019.

Miller MB, Bassler BL. 2001. Quorum Sensing in Bacteria.

Moran NA, McCutcheon JP, Nakabachi A. 2008. Genomics and Evolution of Heritable Bacterial Symbionts. Annual Review of Genetics 42:165–190. DOI: 10.1146/annurev.genet.41.110306.130119.

Mulio SÅ, Zwolińska A, Klejdysz T, Prus-Frankowska M, Michalik A, Kolasa M, Łukasik P. 2024. Limited variation in microbial communities across populations of *Macrosteles* leafhoppers (Hemiptera: Cicadellidae). Evolutionary Biology. DOI: 10.1101/2024.01.28.577611.

Munoz MM, Spencer N, Enomoto S, Dale C, Rio RVM. 2020. Quorum sensing sets the stage for the establishment and vertical transmission of *Sodalis praecaptivus* in tsetse flies. PLOS Genetics 16:e1008992. DOI: 10.1371/journal.pgen.1008992.

Nakabachi A, Inoue H, Hirose Y. 2022. Microbiome analyses of 12 psyllid species of the family Psyllidae identified various bacteria including *Fukatsuia* and *Serratia symbiotica*, known as secondary symbionts of aphids. BMC Microbiology 22:15. DOI: 10.1186/s12866-021-02429-2.

Neuwirth E. 2022. RColorBrewer: ColorBrewer Palettes.

Parada AE, Needham DM, Fuhrman JA. 2016. Every base matters: assessing small subunit rRNA primers for marine microbiomes with mock communities, time series and global field samples: Primers for marine microbiome studies. Environmental Microbiology 18:1403–1414. DOI: 10.1111/1462-2920.13023.

Pinheiro J, Bates D, R Core Team. 2023. nlme: Linear and Nonlinear Mixed Effects Models.

Prodan A, Tremaroli V, Brolin H, Zwinderman AH, Nieuwdorp M, Levin E. 2020. Comparing bioinformatic pipelines for microbial 16S rRNA amplicon sequencing. PLOS ONE 15:e0227434. DOI: 10.1371/journal.pone.0227434.

Props R, Kerckhof F-M, Rubbens P, De Vrieze J, Hernandez Sanabria E, Waegeman W, Monsieurs P, Hammes F, Boon N. 2017. Absolute quantification of microbial taxon abundances. The ISME Journal 11:584–587. DOI: 10.1038/ismej.2016.117.

Quast C, Pruesse E, Yilmaz P, Gerken J, Schweer T, Yarza P, Peplies J, Glöckner FO. 2013. The SILVA ribosomal RNA gene database project: improved data processing and web-based tools. Nucleic Acids Research 41:D590–D596. DOI: 10.1093/nar/gks1219.

R Core Team. 2023. R: A Language and Environment for Statistical Computing.

Ravenscraft A, Berry M, Hammer T, Peay K, Boggs C. 2019. Structure and function of the bacterial and fungal gut microbiota of Neotropical butterflies. Ecological Monographs 89:e01346. DOI: 10.1002/ecm.1346.

Rognes T, Flouri T, Nichols B, Quince C, Mahé F. 2016. VSEARCH: a versatile open source tool for metagenomics. PeerJ 4:e2584. DOI: 10.7717/peerj.2584.

Salter SJ, Cox MJ, Turek EM, Calus ST, Cookson WO, Moffatt MF, Turner P, Parkhill J, Loman NJ, Walker AW. 2014. Reagent and laboratory contamination can critically impact sequence-based microbiome analyses. BMC Biology 12:87. DOI: 10.1186/s12915-014-0087-z.

Sánchez-Bayo F, Wyckhuys KAG. 2019. Worldwide decline of the entomofauna: A review of its drivers. Biological Conservation 232:8–27. DOI: 10.1016/j.biocon.2019.01.020.

Sanders JG, Łukasik P, Frederickson ME, Russell JA, Koga R, Knight R, Pierce NE. 2017. Dramatic differences in gut bacterial densities correlate with diet and habitat in rainforest ants. Integrative and Comparative Biology 57:705–722. DOI: 10.1093/icb/icx088.

Shannon CE. 1948. A mathematical theory of communication. The Bell System Technical Journal 27:379–423, 623–656. DOI: 10.1002/j.1538-7305.1948.tb01338.x.

Sørensen T. 1948. A method of establishing groups of equal amplitude in plant sociology based on similarity of species and its application to analyses of the vegetation on Danish commons. Biol. Skr. 5:1–34.

Srivathsan A, Ang Y, Heraty JM, Hwang WS, Jusoh WFA, Kutty SN, Puniamoorthy J, Yeo D, Roslin T, Meier R. 2023. Convergence of dominance and neglect in flying insect diversity. Nature Ecology & Evolution 7:1012–1021. DOI: 10.1038/s41559-023-02066-0.

Srivathsan A, Feng V, Suárez D, Emerson B, Meier R. 2024. ONTbarcoder 2.0: rapid species discovery and identification with real-time barcoding facilitated by Oxford Nanopore R10.4. Cladistics 40:192–203. DOI: 10.1111/cla.12566.

Srivathsan A, Lee L, Katoh K, Hartop E, Kutty SN, Wong J, Yeo D, Meier R. 2021. ONTbarcoder and MinION barcodes aid biodiversity discovery and identification by everyone, for everyone. BMC Biology 19:217. DOI: 10.1186/s12915-021-01141-x.

Stork NE. 2018. How many species of insects and other terrestrial arthropods are there on earth? Annual Review of Entomology 63:31–45. DOI: 10.1146/annurev-ento-020117-043348.

Surmacz B, Stec D, Prus-Frankowska M, Buczek M, Michalczyk Ł, Łukasik P. 2024. Pinpointing the microbiota of tardigrades: what is really there? :2024.01.24.577024. DOI: 10.1101/2024.01.24.577024.

Tourlousse DM, Yoshiike S, Ohashi A, Matsukura S, Noda N, Sekiguchi Y. 2017. Synthetic spike-in standards for high-throughput 16S rRNA gene amplicon sequencing. Nucleic Acids Research 45:e23. DOI: 10.1093/nar/gkw984.

Truett GE, Heeger P, Mynatt RL, Truett AA, Walker JA, Warman ML. 2000. Preparation of PCR-quality mouse genomic DNA with hot sodium hydroxide and Tris (HotSHOT). BioTechniques 29:52–54. DOI: 10.2144/00291bm09.

Van Klink R, Bowler DE, Gongalsky KB, Swengel AB, Gentile A, Chase JM. 2020. Meta-analysis reveals declines in terrestrial but increases in freshwater insect abundances. Science 368:417–420. DOI: 10.1126/science.aax9931.

Vasilita C, Feng V, Hansen AK, Hartop E, Srivathsan A, Struijk R, Meier R. 2024. Express barcoding with NextGenPCR and MinION for species-level sorting of ecological samples. Molecular Ecology Resources 24:e13922. DOI: 10.1111/1755-0998.13922.

Vázquez-Baeza Y, Pirrung M, Gonzalez A, Knight R. 2013. EMPeror: a tool for visualizing high-throughput microbial community data. GigaScience 2:2047–217X-2–16. DOI: 10.1186/2047-217X-2-16.

Vesterinen EJ, Ruokolainen L, Wahlberg N, Peña C, Roslin T, Laine VN, Vasko V, Sääksjärvi IE, Norrdahl K, Lilley TM. 2016. What you need is what you eat? Prey selection by the bat *Myotis daubentonii*. Molecular Ecology 25:1581–1594. DOI: 10.1111/mec.13564.

Weiss S, Xu ZZ, Peddada S, Amir A, Bittinger K, Gonzalez A, Lozupone C, Zaneveld JR, Vázquez-Baeza Y, Birmingham A, Hyde ER, Knight R. 2017. Normalization and microbial differential abundance strategies depend upon data characteristics. Microbiome 5:27. DOI: 10.1186/s40168-017-0237-y.

Weisser W, Siemann E. 2008. Insects and Ecosystem Function. DOI: 10.1007/978-3-540-74004-9.

Whittaker RH. 1972. Evolution and measurement of species diversity. Taxon 21:213–251. DOI: 10.2307/1218190.

Wickham H. 2016. ggplot2: Elegant Graphics for Data Analysis. Springer-Verlag New York.

Wickham H, François R, Henry L, Müller K, Vaughan D. 2023. dplyr: A Grammar of Data Manipulation.

Wong CNA, Ng P, Douglas AE. 2011. Low-diversity bacterial community in the gut of the fruitfly *Drosophila melanogaster*. Environmental Microbiology 13:1889–1900. DOI: 10.1111/j.1462-2920.2011.02511.x.

Zhang J, Kobert K, Flouri T, Stamatakis A. 2014. PEAR: a fast and accurate Illumina Paired-End reAd mergeR. Bioinformatics 30:614–620. DOI: 10.1093/bioinformatics/btt593.

