## Supplementary figures and images for "Implementing high-throughput insect barcoding in microbiome studies: impact of non-destructive DNA extraction on microbiome reconstruction"

### Suppl. Figure S1

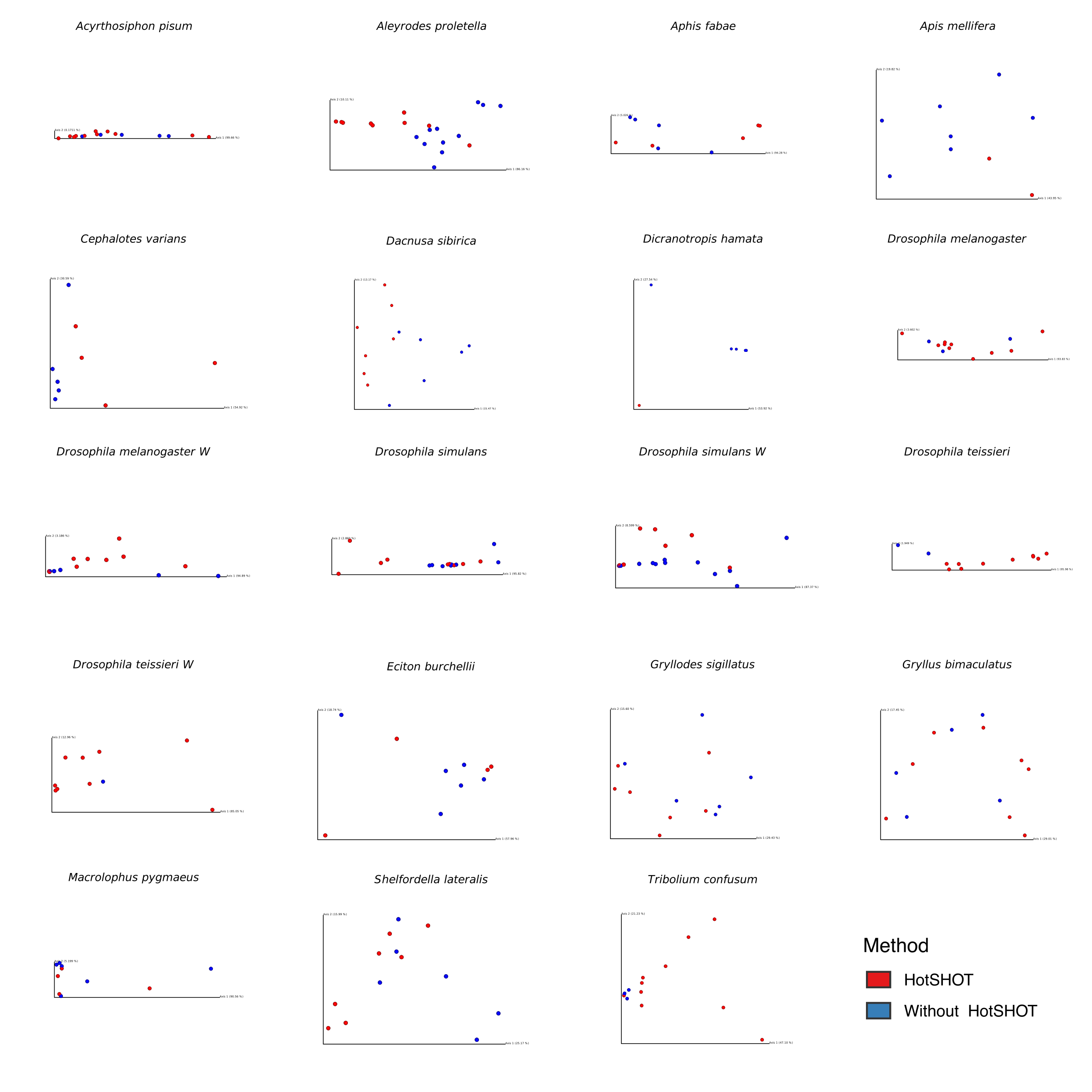
